# Control of MYC-dependent apoptotic threshold by a co-amplified ubiquitin E3 ligase UBR5

**DOI:** 10.1101/515247

**Authors:** Xi Qiao, Ying Liu, Maria Llamazares Prada, Abhishekh Gupta, Alok Jaiswal, Mukund Sharma, Heidi Haikala, Kati Talvinen, Laxman Yetukuri, Joanna W. Pylvänäinen, Juha Klefström, Pauliina Kronqvist, Annika Meinander, Tero Aittokallio, Ville Hietakangas, Martin Eilers, Jukka Westermarck

## Abstract

MYC protein expression has to be tightly controlled to allow for maximal cell proliferation without inducing apoptosis. Here we discover UBR5 as a novel MYC ubiquitin ligase and demonstrate how it functions as a molecular rheostat to prevent excess accumulation of MYC protein. UBR5 effects on MYC protein stability are independent on N-terminal FBW7 degron of MYC. Endogenous UBR5 inhibition induces MYC protein expression and activates MYC target genes. Moreover, UBR5 governs MYC-dependent phenotypes *in vivo* in *Drosophila*. In cancer cells, UBR5-mediated MYC protein suppression diminishes cell killing activity of cancer therapeutics. Further, we demonstrate that UBR5 dominates MYC protein expression at the single-cell level in human basal-type breast cancer tissue. *Myc* and *Ubr5* are co-amplified in MYC-driven human cancer types, and UBR5 controls MYC-mediated apoptotic threshold in co-amplified basal type breast cancer cells. In summary, UBR5 is a novel MYC ubiquitin ligase and an endogenous rheostat for MYC protein expression *in vivo*. Clinically, expression of UBR5 may be important for protection of breast cancer cells from drug-induced, and MYC-dependent, apoptosis.

## Introduction

Emerging notion of overall poor concordance between gene amplifications and corresponding protein expression levels in cancer (Mertins, Mani et al., 2016) highlights the need for better understanding of how protein expression levels are regulated at post-genomic level. Transcription factor MYC regulates numerous physiological and pathological processes (Dang, 2012, Grifoni & Bellosta, 2015, McMahon, 2014). Critically, distinct protein expression thresholds dictate MYC’s biological output *in vitro* and *in vivo* (McMahon, 2014, Murphy, Junttila et al., 2008, Nieminen, Eskelinen et al., 2013, Sarosiek, Fraser et al., 2017). In cancers, MYC is a classic example of an oncoprotein whose overexpression at protein level do not match neither with the extent of mRNA overexpression, nor with the frequency of genetic amplifications (Farrell & Sears, 2014, Levens, 2010, Xu, Chen et al., 2010). Therefore, a deeper understanding of endogenous mechanisms that govern MYC protein expression levels, and MYC-induced phenotypes, in normal and diseased tissues is of great general relevance.

MYC mRNA transcription, and MYC protein activity are both required for proliferation induction in several types of normal cells (Dang, 2012, Myant, Qiao et al., 2015, Trumpp, Refaeli et al., 2001). In addition to cell culture and mouse models (Murphy et al., 2008, Myant et al., 2015, Pelengaris, Khan et al., 2002, Sarosiek et al., 2017, Trumpp et al., 2001), the physiological role of MYC has been widely studied in *Drosophila*, where the conserved homolog of MYC, dMyc regulates tissue growth and animal size (Grifoni & Bellosta, 2015). In particular, overexpression of dMyc induces growth of cells of the imaginal discs, which are the organs giving rise to wings in adult fly (Grifoni & Bellosta, 2015). MYC is also a critical driver of malignant growth of human cancer cells (Dang, 2012). However, MYC protein levels exceeding the optimal proliferation promoting levels sensitizes both normal and cancer cells to apoptosis induction (apoptosis priming)(McMahon, 2014, Murphy et al., 2008, Nieminen et al., 2013, Pelengaris et al., 2002). This principle was originally revealed by findings that highly overexpressed transgenic MYC initiates both proliferation and apoptosis programs in pancreatic islet cells, but MYC overexpression was capable to drive tumourigenesis only in the context of efficient apoptosis suppression (Pelengaris et al., 2002). More recently, the *in vivo* importance of MYC expression thresholds in defining the balance between proliferation and apoptosis was validated by allelic series of inducible MYC expression (Murphy et al., 2008). In addition to spontaneous apoptosis, high MYC levels prime tumour cells to apoptosis induction by drugs that interfere with high replication and cell division rates, such as topoisomerase inhibitors or taxanes (McMahon, 2014, Rohban & Campaner, 2015, Topham, Tighe et al., 2015). Indeed, MYC-mediated mitochondrial apoptosis regulation governs both developmental and cancer-therapy induced apoptosis *in vivo* (Sarosiek et al., 2017). Thus, MYC protein expression thresholds control the balance between proliferation and apoptosis priming both in normal and cancer cells. However, how MYC protein levels are controlled endogenously to maintain an optimal MYC balance is incompletely understood.

UBR5 (Ubiquitin protein ligase E3 component n-recognin 5) is an evolutionary conserved E3 ubiquitin ligase that destabilize proteins with N-terminal recognition sequences exposed by proteolytic cleavage (Shearer, Iconomou et al., 2015, Varshavsky, 2011). Recently UBR5 has been shown to target substrates also through other recognition mechanisms than N-terminal recognition (Shen, Qiu et al., 2018). UBR5 is essential for mammalian development(Saunders, Hird et al., 2004), and has been linked to both pro-tumourigenic and tumour suppressor activities (Clancy, Henderson et al., 2003, Liao, Song et al., 2017, Saunders et al., 2004). However, the mechanism by which UBR5 would promote tumour growth, and how the pro-tumourigenic effects of UBR5 in human cancer cells(Clancy et al., 2003, Liao et al., 2017), can be consolidated with its suggested role as a tumour suppressor in Drosophila (Clancy et al., 2003, Saunders et al., 2004), remains as critical unanswered questions (Shearer et al., 2015).

Here, we have identified UBR5 as a ubiquitin ligase for MYC. In normal tissues *in vivo*, UBR5 inhibits ectopic growth by suppression of MYC protein expression, whereas in cancer cells UBR5-mediated MYC suppression protects the cells from apoptosis priming. Further, our data reveal genetic co-amplification of *Ubr5* and *Myc* in several solid human cancers; and demonstrate functional relevance of reciprocal protein level regulation between these two widely amplified cancer genes in defining the apoptosis sensitivity of breast cancer cells. Attenuation of MYC protein levels below the apoptosis priming threshold thereby provides a plausible explanation for pro-tumourigenic function of UBR5 in cancers. We further demonstrate that UBR5 inhibition shifts cancer cell populations towards MYC^high^ status that primes them for efficient cell killing by clinically used cancer therapies. Thereby, these results may also provide an important cue for development of therapeutic strategies for targeting tumour tissues by increasing the apoptosis priming function of MYC.

## Results

### UBR5 suppress MYC protein expression

We conducted a screen for ubiquitin ligases that regulate MYC protein abundance with siRNAs against 591 ubiquitin ligases (Fig. 1a). To exclude ubiquitin ligases that regulate MYC stability via the well-characterized MYC ubiquitin ligase FBW7 (Welcker & Clurman, 2008), the siRNA library was screened against U2OS cells stably expressing MYCT58A mutant (threonine 58 mutated to alanine), that is resistant to FBW7-mediated destabilization (Welcker, Orian et al., 2004). 48 hours after siRNA transfection, cells were treated for 3,5 hours with cycloheximide to emphasize the impact of protein stability regulation by siRNA library targets, and immunofluorescence (IF) detection of MYC was thereafter used as a read-out in a high-content imaging-based assay (Fig. 1a). Among a small group of siRNAs consistently affecting MYCT58A levels (Table S1), HECT-domain containing E3 ligase UBR5 (alias EDD)(Shearer et al., 2015) was the only one that did not affect MYC mRNA expression (Fig. S1a), and was therefore selected for further validation experiments.

**Fig. 1.**
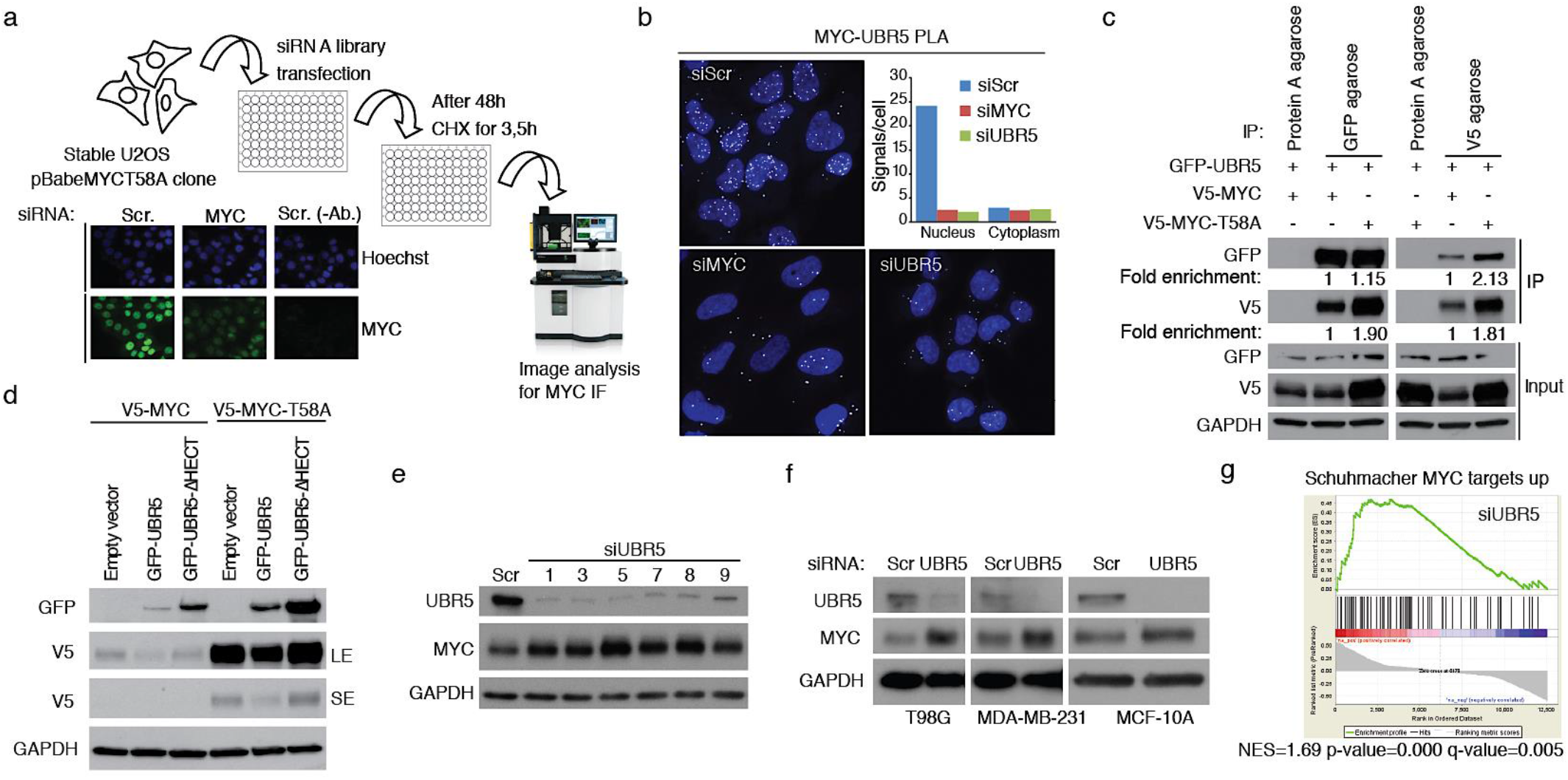
UBR5 negatively regulates MYC protein expression. **a**, High-content imaging-based screen was performed to identify E3 ubiquitin ligases that regulate MYCT58A. U2OS cells stably expressing MYCT58A mutant were transfected with siRNAs against 591 E3 ubiquitin ligases. After 48 hours from siRNA transfection, cells were treated with cycloheximide. MYC (N262) antibody was used to detect MYC expression. An example of immunofluorescence staining of MYC and a secondary antibody control only is shown. **b**, Proximity ligation assay analysis of the UBR5 and MYC association in HeLa cells transfected with scrambled (Scr), MYC siRNA or UBR5 siRNA. Shown is a quantification of the average number of UBR5-MYC PLA signals per cell in cytoplasm and nucleus. **c**, UBR5, wild-type MYC or MYCT58A mutants were overexpressed in HEK293 cells by transient transfection. Cell lysates were immunoprecipitated with GFP or V5 agarose and interaction was detected with relevant antibodies. The numbers show fold change of relevant proteins. Shown is a representative experiment; n=3. **d**, HEK293 cells were cotransfected with empty vector, GFP-tagged UBR5, UBR5 mutant, V5-tagged MYC and MYCT58A mutant plasmids. After 48 hours transfection, the cell lysates were collected for immunoblotting. Shown is a representative experiment; n=2. **e**, Hela cells were transfected with scrambled (Scr) or 6 different UBR5 siRNAs. After 72 hours, cell lysates were collected to detect MYC expression by western blot. **f**, T98G, MDA-MB-231 and MCF10A cells were transfected with scrambled (Scr) or UBR5 (5#) siRNA. After 72 hours, cells lysates were collected to detect MYC expression. **g**, Gene set enrichment analysis (GSEA) of RNA-seq data generated in Hela cells transfected with Scr or UBR5 siRNAs was performed by employing gene sets from the Molecular Signature Database (MSigDB). GSEA analysis demonstrated that MYC-induced genes were enriched upon UBR5 inhibition. NES, normalized enrichment score.

Physical association between endogenous UBR5 and MYC in HeLa cell nuclei was confirmed by proximity ligation analysis (PLA)(Fig. 1b). The specificity of the PLA reaction was confirmed by staining with PLA secondary antibodies alone (Fig. S1b), and by a clear decrease of positive MYC-UBR5 PLA signals, and of UBR5 immunofluorescence signal by siRNA treatments (Fig. 1b and Fig. S1c,d). Whereas MYC and UBR5 siRNA treatments exclusively decreased the number of nuclear PLA signals, they did not affect cytoplasmic signals (Fig. 1b). Therefore, we conclude that the sparse cytoplasmic PLA signals represent staining background and that MYC-UBR5 association occurs in the nucleus.

To confirm FBW7-independent function of UBR5, we studied whether MYC binding to UBR5 is affected by MYC T58A mutation, that abolishes recognition by FBW7 (Farrell & Sears, 2014, Gregory, Qi et al., 2003, Welcker et al., 2004). As expected, V5-MYC-T58A was expressed at higher level than wild-type V5-MYC, but in reciprocal co-immunoprecipitation analysis, GFP-UBR5 associated equally efficiently with both forms of MYC (Fig. 1c). By using different MYC fragments, we delineated aminoacids 181-262 of MYC as the minimal common region for MYC-UBR5 association (Fig. S1e). This data further indicates that the aminoterminal T58 phosphorylation site is not necessary MYC-UBR5 interaction. Additionally, UBR5 overexpression suppressed protein expression of MYC T58A to a similar degree as that of WT MYC (Fig. 1d). Importantly, UBR5 mutant lacking the ubiquitin ligase HECT domain was not able to suppress MYC expression (Fig. 1d). This supports the expected role of UBR5 in regulating MYC via its ubiquitin ligase function.

The effects of UBR5 depletion on endogenous MYC protein levels in HeLa cells was confirmed by immunoblot analysis using six independent UBR5 siRNA sequences (Fig. 1e). In addition to HeLa cells, endogenous MYC protein induction by UBR5 inhibition was confirmed in T98G glioblastoma, and MDA-MB-231 breast cancer cells, and in immortalized MCF-10A mammary epithelial cells (Fig. 1f). This demonstrates that the function of UBR5 as a negative regulator of MYC expression is neither lineage-specific, nor restricted to transformed cells. In addition, we performed RNA-sequencing analysis of HeLa cells depleted of MYC or UBR5 (Fig. S1f). By gene set enrichment analysis (GSEA), we found that UBR5 depletion induces expression of same MYC target signature (Fig. 1g) that is suppressed by MYC depletion (Fig. S1g).

In summary, these experiments identify UBR5 as a novel FBW7, and threonine 58 phosphorylation, -independent regulator of MYC protein expression and activity.

### UBR5 is a MYC ubiquitin ligase

To validate that UBR5-mediated MYC regulation occurs at the protein level, we compared the effect of UBR5 depletion on MYC mRNA and protein expression in HeLa cells. Based on quantifications from a number of independent repeat experiments, UBR5 depletion induced a robust MYC protein induction, whereas its impact on the MYC mRNA expression was minimal (Fig. 2a). Further supporting regulation at post-transcriptional levels, UBR5 depletion significantly potentiated MYC protein expression when ectopic MYC mRNA expression was induced by gradual removal of doxycycline from mouse OS-Tet-Off-MYC cells(Jain, Arvanitis et al., 2002) (Fig. 2b). Consistently with UBR5 regulating MYC protein stability, UBR5 depletion stabilized MYC in a cycloheximide chase experiment in HeLa (Fig. 2c and S2a), and in U2OS cells (Fig. S2b,c). Furthermore, depletion of either FBW7 (Fig. S2d) or UBR5 increased endogenous MYC stability, and co-depletion of both ubiquitin ligases resulted in synergistic stabilization (Fig. 2d and S2e). In addition to demonstrating that UBR5 regulates MYC stability, these results provide yet another evidence for independent roles of UBR5 and FBW7 in MYC protein regulation. Consistent with ubiquitination-mediated regulation of MYC by UBR5 (Fig. 1d), MYC inhibition by overexpression of the wild-type UBR5 was rescued by co-treatment with proteasome inhibitor MG132 (Fig. 2e, f). Moreover, in ubiquitination assays, only overexpression of the wild-type UBR5, but not the HECT mutant, induced ubiquitination of V5-MYC (Fig. 2g). Reciprocally, siRNA-mediated depletion of UBR5 potently inhibited MYC ubiquitination (Fig. 2h). Moreover, we found that UBR5-induced ubiquitination of MYC is via K48-linked ubiquitin chains (Fig. 2i) which are mainly responsible for proteasomal degradation of proteins.

**Fig. 2.**
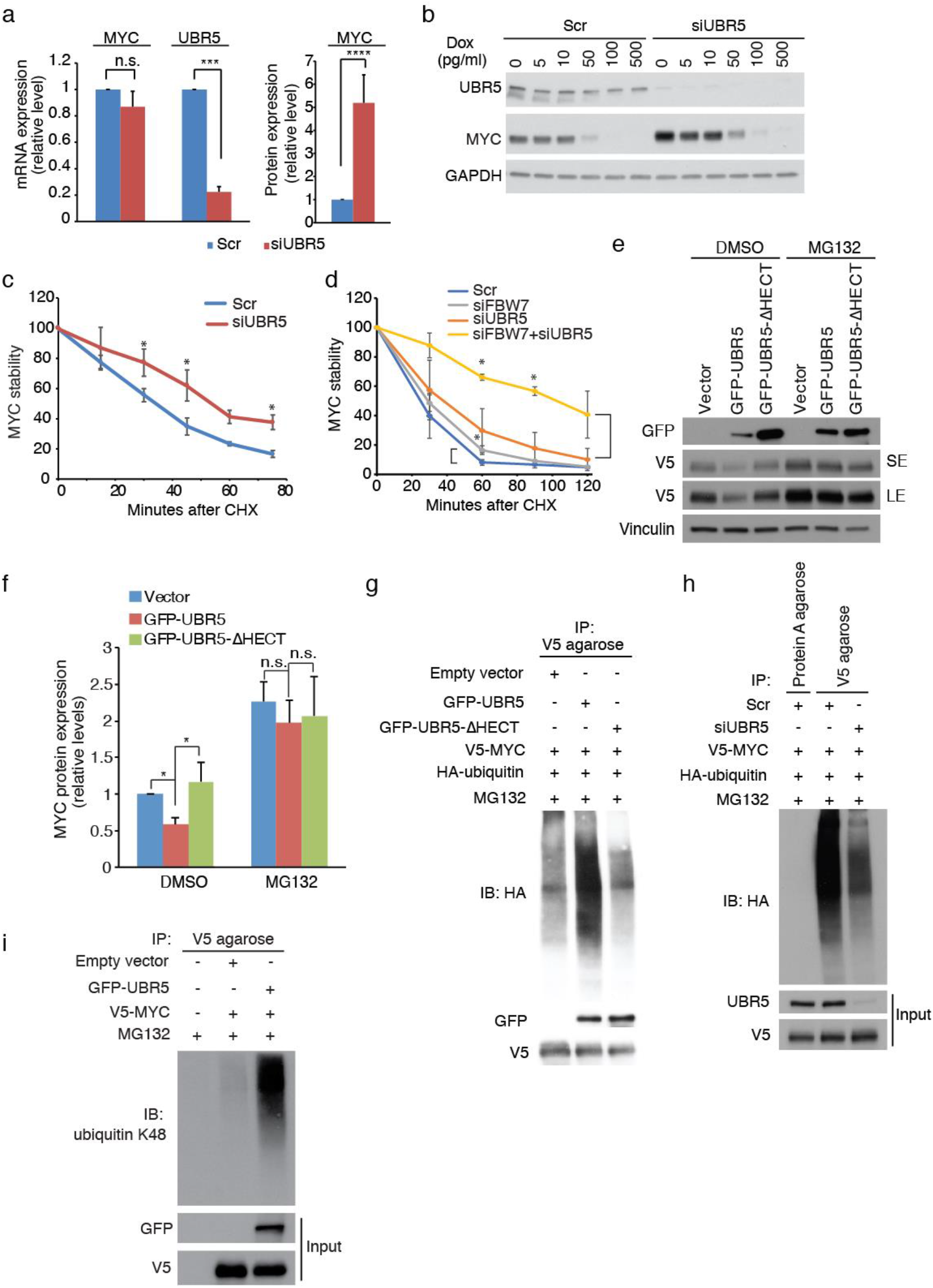
Identification of UBR5 as a MYC ubiquitin ligase. **a**, Analysis of MYC mRNA and protein expression in Hela cells transfected with scrambled (Scr) or UBR5 (5#) siRNA. Error bars represent SD (for mRNA expression, n=3; for protein expression, n=14), ***, P < 0.001, ****, P < 0.0001 by student’s t-test. **b**, Western blot analysis of MYC expression in osteosarcoma-MYC-off cell line derived from transgenic mice. The cells cultured with 20ng/ml doxycycline were transfected with scrambled (Scr) or UBR5 siRNA. After 48 hours culture in medium with 20ng/ml doxycycline, the medium was changed with indicated concentrations of doxycycline for another 48 hours. The total cell lysates were subjected to immunoblotting with indicated antibodies. **c**, Hela cells transfected with scrambled (Scr) or UBR5 (5#) siRNA for 72 hours were treated with cycloheximide (CHX, 60μg/ml) for indicated time points. Cell lysates were examined for MYC expression by western blot (see Fig S2a). Error bars mean ± SD. *, P < 0.05 by student’s t-test (n=3). **d**, Hela cells transfected with indicated siRNAs for 72 hours were treated with cycloheximide (CHX, 60μg/ml) for different time points. Cell lysates were examined with MYC antibody by western blot (see Fig S2e). Error bars mean ± SD. *, P < 0.05 by student’s t-test (n=3). **e**, HEK293 cells were co-transfected with GFP-tagged wild type UBR5, UBR5-HECT domain mutant, and V5-tagged MYC plasmid. After 48 hours transfection, the cells were treated with MG132 (20μM) 6 hours. The cell lysates were analyzed by immunoblotting. SE and LE represent short and long exposure time. **f**, Quantification of MYC expression from (e). Error bars show SD from 4 independent experiments. *, P < 0.05 by student’s t-test. **g**, HEK293 cells were co-transfected with indicated plasmids. After 48 hours transfection, the cells were treated with MG132 (20μM) for 6 hours. Immunoprecipitation was performed with anti-V5 agarose. Ubiquitination of MYC was detected by HA antibody. **h**, Hela cells were co-transfected as Scr siRNA, UBR5 siRNA and indicated plasmids. The cells were treated with MG132 (20μM) for 6 hours. Immunoprecipitation was performed with anti-V5 agarose. Ubiquitination of MYC was detected by HA antibody. **i**, Hela cells were transfected with indicated plasmids. The cells were treated with MG132 (20μM) for 6 hours. Immunoprecipitation was performed with V5 agarose. Ubiquitination of MYC was detected by ubiquitin K48 specific antibody.

Together, these data demonstrate that MYC regulation by UBR5 occurs at the protein level and identify UBR5 as a MYC ubiquitin ligase.

### UBR5 controls *in vivo* tissue growth in dMyc-dependent manner in *Drosophila*

To explore the functional role of the UBR5-MYC axis in normal growth, we utilized a *Drosophila melanogaster* model in which UBR5 ortholog Hyd (Hyperplastic discs) was originally identified as a growth suppressor(Clancy et al., 2003, Mansfield, Hersperger et al., 1994, Shearer et al., 2015). RNAi-mediated depletion of Hyd in *Drosophila* S2 cells (Fig. S3a) led to elevated levels of *Drosophila* MYC (dMyc) protein (Fig. 3a), whereas the mRNA levels of dMyc did not correlate with dMYC protein inudction (Fig. 3b). Importantly, the levels of Hyd RNA reduction by RNAi was directly reflected in a comparable 2-fold induction of dMyc protein expression (Fig. S3a and 3a). Similar to regulation of MYC targets by UBR5 in human cells (Fig. 1g), Hyd depletion induced mRNA expression of two dMyc targets eIF6 and Nop5 (Fig. 3c and S3b), and protein expression of Fibrillarin (Fig. 3d,e), which is a well-established MYC target in both human and *Drosophila*.

**Fig. 3.**
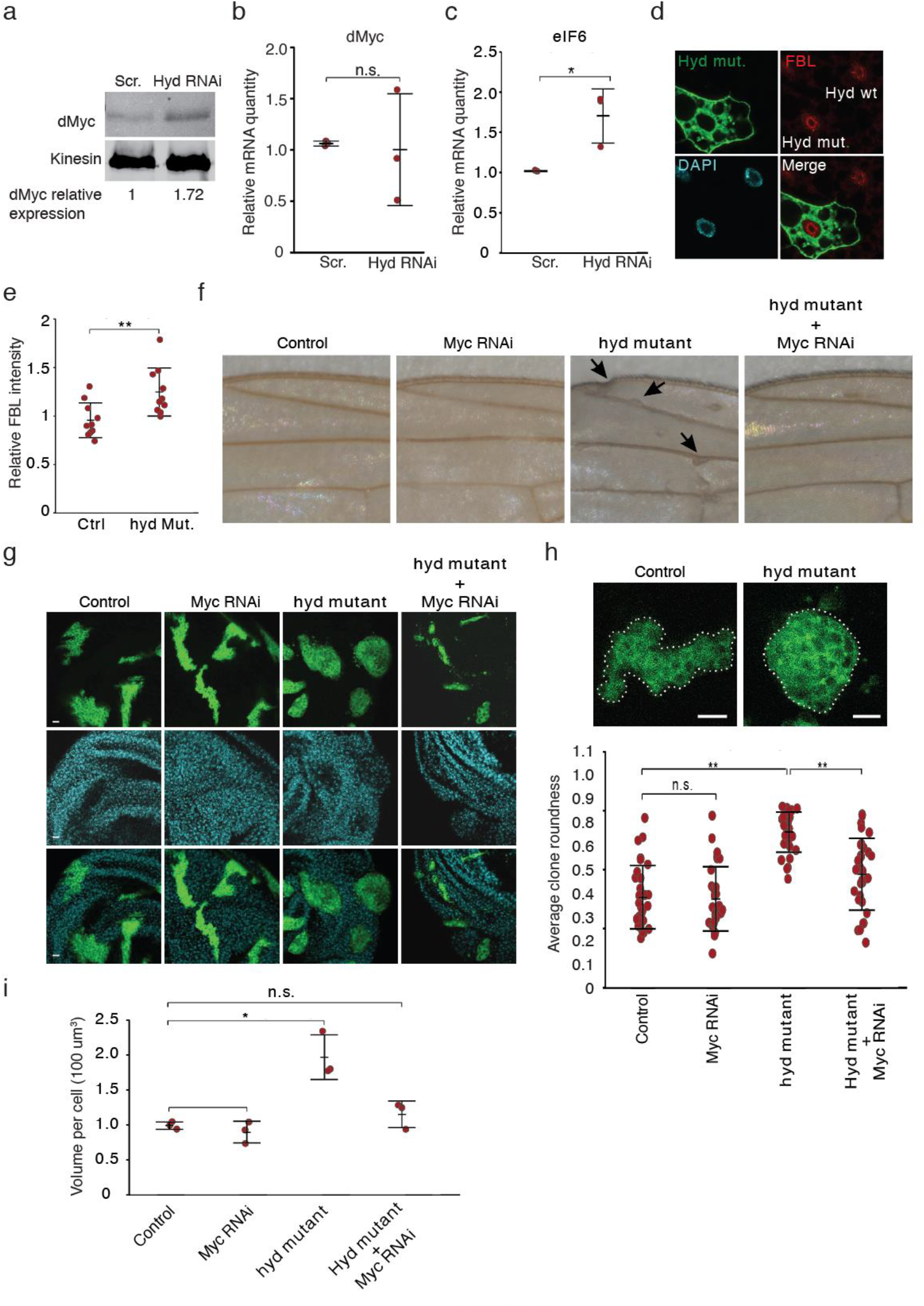
Drosophila UBR5 (Hyd) inhibits wing imaginal disc overgrowth in Myc-dependent manner. **a**, Immunoblot showing immunoprecipitated dMyc levels in S2 cells transfected with Scr or Hyd dsRNA. Kinesin was used as a control for the input. Shown is a representative blot of two experiments with similar results. **b**, Quantitative RT-PCR analysis of dMyc mRNA expression in control and Hyd depleted S2 cells. RP49 was used as a reference gene. Error bar means SD. n=3. **c**, Quantitative RT-PCR analysis of Myc target gene, eIF6 mRNA expression in control and Hyd depleted S2 cells. Error bar means SD. n=3. *, P < 0.05 by student’s t-test. **d**, Loss of Hyd in GFP-labelled fat body mutant clones leads to increase of Fibrillarin. **e**, Quantification of Fig. 3d. The Fibrillarin level of an adjacent GFP negative cell was used as reference. **, P < 0.01 by student’s t-test. **f**, Induction of Hyd mutant clones leads to uneven wing morphology, which in suppressed by simultaneous knockdown of Myc within the clones. Arrows in magnified image show uneven wing margins and veins. **g**, Loss of Hyd in GFP-labelled mutant clones leads to overgrowth indicated by roundness, and loss of normal epithelial morphology. Simultaneous knockdown of dMyc strongly reduces the roundness of the Hyd mutant clones. Scale bar means 10μm. **h**, Representative images show clone morphology in wing imaginal discs in control and hyd mutant clones. Scale bar means 10μm. Quantification of the clone roundness was shown. Error bar means SD. n=30. **, P <0.01 by student’s t-test. **i**, Loss of Hyd in GFP-labelled wing disc mutant clones leads to increased cell volume. Simultaneous knockdown of dMyc fully suppresses this phenotype. *, P < 0.05 by student’s t-test.

*Drosophila* models allow the use of somatic recombination to generate clones of mutant tissue in an otherwise heterozygous background. Clones of *hyd*^K3.5^ mutant generated during mid larval development (72 h after egg laying) led to a phenotype clearly visible in the adult wings (Fig. S3c). Wings with *hyd* mutant clones were irregular, lacking the normal flat wing morphology, and displaying uneven wing margins and veins (Fig. 3f and S3c, see arrows). Interestingly, this irregular wing phenotype was strongly suppressed by simultaneous knockdown of dMyc in the mutant clones (Fig. 3f and S3c).

The use of the MARCM system to generate mutant clones allowed us to GFP label, and to visualize the tissue morphology of the wing imaginal discs, the larval wing precursors (Lee & Luo, 1999). Control and Myc RNAi clones appeared morphologically normal with GFP positive clones having a normal but irregular shape (Fig. 3g,h). In contrast, *hyd* mutant clones had a round morphology with clusters of cells growing out from the epithelial plane (Fig. 3g,h), which was confirmed by quantification of the clone roundness (Fig. 3h). The roundness phenotype in wing clones has been earlier validated to mark activated Ras-MAPK-MYC signalling (Prober & Edgar, 2002). The large round *hyd* mutant clones were present in all regions of the imaginal disc, which is consistent with our findings of morphological irregularities throughout the adult wing. Most importantly, simultaneous RNAi-mediated knockdown of dMyc suppressed the roundness morphology of the *hyd* mutant clones, which was further confirmed by quantification (Fig. 3h).

dMyc has an important role in regulation of *Drosophila* cell size(Trumpp et al., 2001). To dissect the relevance of cell size and proliferation in dMyc-dependent regulation of *Drosophila* tissue growth by Hyd, we first analyzed cell size in *hyd* mutant clones. *hyd* mutant cells were significantly larger than controls and this phenotype was fully suppressed by simultaneous knockdown of dMyc (Fig. 3i). We also observed higher percentage of phospho-histone H3, a marker commonly used as a proxy for cell proliferation, in the *hyd* mutant clones, but this phenotype was independent of dMyc expression (Fig. S3d). Thus, we conclude that the increased cell size, and roundness (established mark of activated MYC signaling (Prober & Edgar, 2002)), but not proliferation, are dependent on dMyc in *hyd* mutant *Drosophila* tissues.

In conclusion, these results demonstrate that in *Drosophila* suppression of dMyc protein expression is a critical part of growth control by the ortholog of UBR5 *in vivo*. Moreover, together with the results from mouse osteosarcoma cells (Figure 2b), these results also demonstrate that the function of UBR5 as a suppressor of MYC is conserved between at least three species.

### UBR5 suppresses MYC-mediated apoptosis in cancer cells

Increased MYC protein expression in cancer cells could lead to either increased proliferation or to apoptosis priming depending on expression levels of MYC (Murphy et al., 2008, Nieminen et al., 2013, Pelengaris et al., 2002, Sarosiek et al., 2017, Topham et al., 2015). In HeLa cells, UBR5 depletion decreased colony growth (Fig. 4a), suggesting that deregulated MYC levels in UBR5-depleted cancer cells may exceed the threshold that lead to apoptosis priming. This was confirmed by the increased PARP cleavage, and by the induction of Caspase-3/7 activity in UBR5-depleted cells (Fig. 4b,c). Interestingly, UBR5 depleted cells with high MYC expression also displayed inhibition of the direct anti-proliferative MYC target gene p21(Seoane, Le et al., 2002) (Fig. 4b). These results indicate that both proliferation and apoptosis priming are initiated upon increased MYC expression, and that loss of cells in colony growth assay is a result of MYC expression levels exceeding the apoptosis threshold. In order to further study whether MYC-induced apoptosis and proliferation are governed by different MYC threshold levels, we titrated UBR5 siRNA amounts and studied the dose-dependent correlation between UBR5 inhibition and expression of MYC, cleaved-PARP, and p21. Interestingly, while p21 suppression was almost maximal already with 10 nM of UBR siRNA (Fig. 4d,e), cleaved-PARP continued to increase with higher UBR5 siRNA concentrations (Fig. 4d,e). Notably, robust PARP induction correlated with induction of MYC expression over two folds threshold level, that was also previously shown to be a threshold level for efficient apoptosis induction by MYC *in vivo* (Murphy et al., 2008). As *in vivo* results also support the conclusion that proliferation induction by MYC requires lower MYC levels than apoptosis induction (Murphy et al., 2008), we propose that that UBR5 has a role in ensuring that MYC remains at proliferation supporting levels and do not exceed the apoptosis priming threshold (Fig. 7h).

**Fig. 4.**
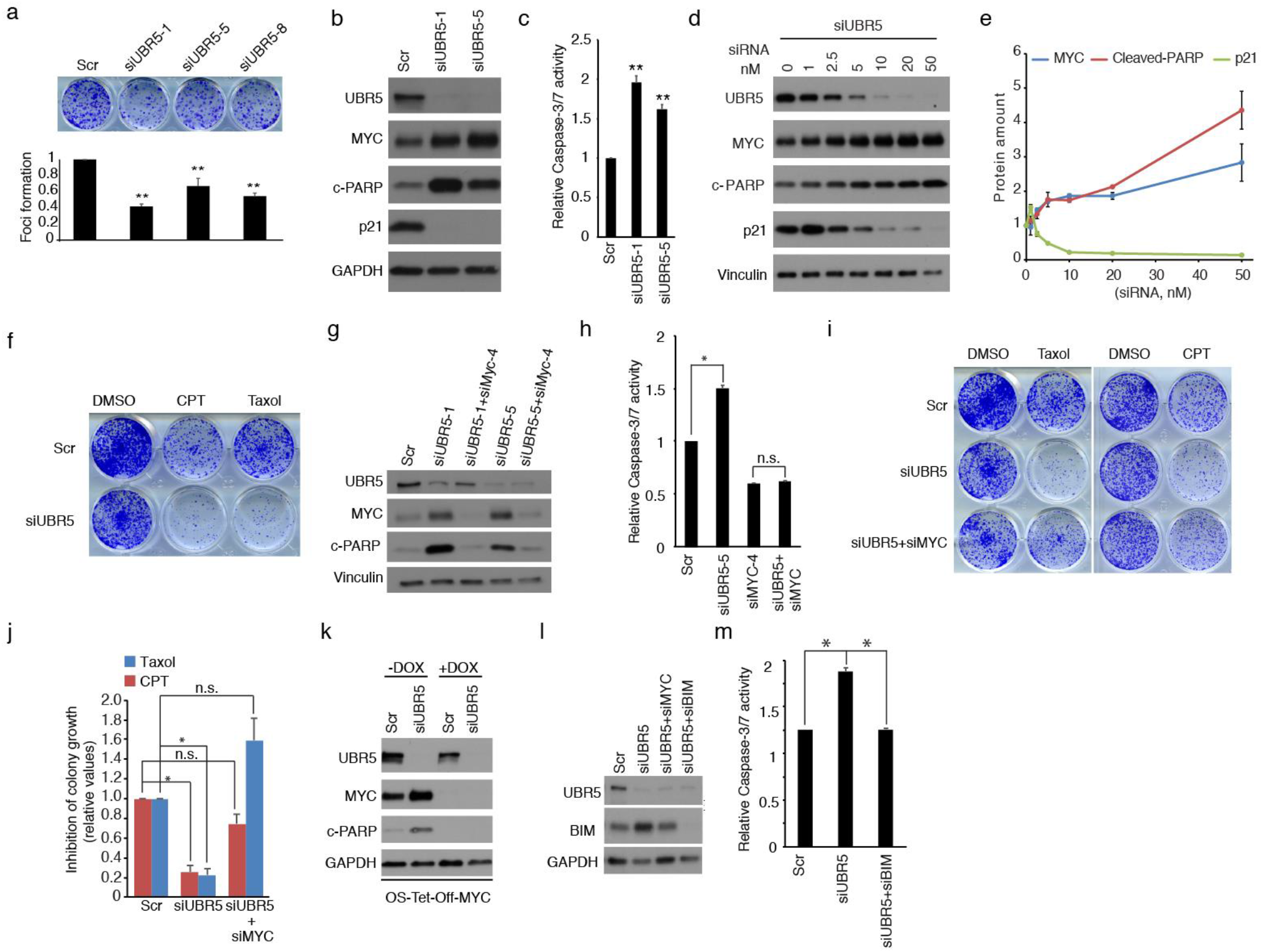
UBR5 suppresses MYC-mediated apoptosis in cancer cells. **a**, Hela cells transfected with three UBR5 siRNAs were cultured 7 days for colony growth assay. Error bars show SD. n=3. **, P <0.01 by student’s t-test. **b**, Hela cells were transfected with two UBR5 siRNAs for 72 hours. Total cell lysates were subjected to immunoblotting with indicated antibodies. **c**, Caspase 3/7 activity was examined in UBR5 depleted Hela cells. Error bars show SD for 7 technical repeats with similar results. **, P <0.01 by student’s t-test. **d**, Hela cells were transfected with different concentrations of UBR5 siRNA for 72 hours. Cell lysates were analysed by western blot with indicated antibodies. **e**, Quantification of MYC, cleared-PARP and p21 expression from (d). Error bars show SD from 2 independent experiments. **f**, Colony growth assay were performed in Hela cells transfected with Scr or siUBR5 siRNA followed by 24 hours treatment of CPT (20nM) or Taxol (1nM). **g**, Hela cells were transfected with two UBR5 siRNAs, or with combination of UBR5 siRNA and MYC siRNA for 72 hours. Cell lysates were collected for western blot with indicated antibodies. **h**, Caspase 3/7 activity was examined in Hela cells transfected with indicated siRNAs. Error bars show SD of 2 independent experiments. *, P < 0.05 by student’s t-test. **i**, Colony growth assay were performed in Hela cells transfected with Scr, siUBR5 or combination of UBR5 and MYC siRNAs followed by 24 hours treatment of Taxol (2nM) or CPT (20nM). After 7 days, cells were fixed and stained. **j**, Quantification of drug response fold changes relative to Scr siRNA in (i). Error bars show SEM of 4 independent experiments. *, P < 0.05 by student’s t-test. **k**, The osteosarcoma-MYC-off cells treated with 20ng/ml doxycycline were transfected with scrambled (Scr) or UBR5 siRNA and cultured in medium with 20ng/ml doxycycline. After 48 hours, medium was changed with or without 20ng/ml doxycycline for culture another 48 hours. Cell lysates were subjected to immunoblotting with indicated antibodies. **l**, Hela cells were transfected with Scr, UBR5 siRNAs, combination of UBR5 and MYC siRNA, and combination of UBR5 and BIM siRNAs. Total cell lysates were subjected to immunoblotting with indicated antibodies. **m**, Caspase 3/7 activity was examined in Hela cells transfected with indicated siRNAs. Shown is mean ± SD of 2 independent experiments. *, P < 0.05 by the student’s t-test.

Apoptosis priming is particularly relevant for increased sensitivity of high MYC expressing cancer cells to antimitotic agents such as taxanes (Topham et al., 2015), and to drugs that interfere with high replication activity, such as topoisomerase I inhibitors (camptothecins) (Rohban & Campaner, 2015). Accordingly, growth inhibition in UBR5 depleted cells was greatly potentiated with treatment of cells with either taxol or camptothecin treatment, at doses that alone do not induce massive loss of cells (Fig 4f). Importantly, both PARP cleavage, and caspase-3/7 activation by UBR5 depletion alone were fully rescued by concurrent MYC depletion (Fig. 4g,h). MYC expression was essential also for hypersensitivity of UBR5 depleted cells to both taxol and camptothecin (Fig. 4i,j). Notably, when considering the degree of drug-induced inhibition of colony growth, the drug responses were indistinguishable between scrambled, and UBR5+MYC siRNA treated cells; and with increasing doses of both drugs (Fig. S4a,b). The important role of MYC in mediating apoptosis priming in cells with low UBR5 expression was further confirmed in OS-Tet-Off-MYC cells. In cells with maximal doxycycline-elicited MYC suppression, UBR5 depletion was unable to induce PARP cleavage (Fig. 4k).

To mechanistically link MYC to the apoptosis sensitisation by UBR5 depletion, we studied the regulation of the pro-apoptotic BH3 protein BIM. BIM was recently shown to be the primary mediator of MYC-induced apoptosis *in vivo* (Muthalagu, Junttila et al., 2014). Accordingly, BIM induction by UBR5 siRNA was abolished by co-depletion of MYC with UBR5 (Fig. 4l), and co-depletion of BIM with UBR5 abolished the induction of apoptosis in UBR5 depleted cells (Fig. 4m). On the other hand, UBR5 depletion inhibited p53, and further depletion of p53 was unable to rescue decreased colony growth or increased PARP cleavage that was induced by UBR5 siRNA (Fig S4c,d), indicating that apoptosis by UBR5 depletion was not mediated by p53.

Together these results demonstrate that UBR5 controls MYC-mediated apoptosis priming and thereby promotes resistance to therapeutic agents frequently used in cancer treatments.

### Co-amplification of *Ubr5* and *Myc* in human breast cancer

The results above suggest that expression of UBR5 might provide a selective advantage for cancer cells by suppressing MYC-mediated apoptosis priming. The cancer types in which this biology is particularly relevant would be expected to exhibit co-expression of *Ubr5* and *Myc* at mRNA level, but UBR5^high^/MYC^low^ status at protein level (see cartoon at Fig. 5a). To identify candidate cancer types with statistically significant (p < 0.05) positive correlation between *Ubr5* and *Myc* mRNA expression, we utilized published gene expression data from 672 cell lines and over 20 different cancer types (Klijn, Durinck et al., 2015). Across all the cancer cell lines, a weak but statistically significant correlation between *Ubr5* and *Myc* mRNA expression levels was detected (Pearson correlation 0.21, p<0.01; Fig. 5b). Among the individual cancer types, significant positive correlation between *Myc* and *Ubr5* mRNA expression was observed in four of the major cancer types, ovary, lymphoid, breast and pancreas (Fig. 5b). Importantly, MYC has been closely linked to progression of all these four cancer types, and existing data indicate a growth-promoting role of UBR5 in at least pancreatic, ovary, and breast cancer (Clancy et al., 2003, Liao et al., 2017, Mann, Ward et al., 2012, O’Brien, Davies et al., 2008).

**Fig. 5.**
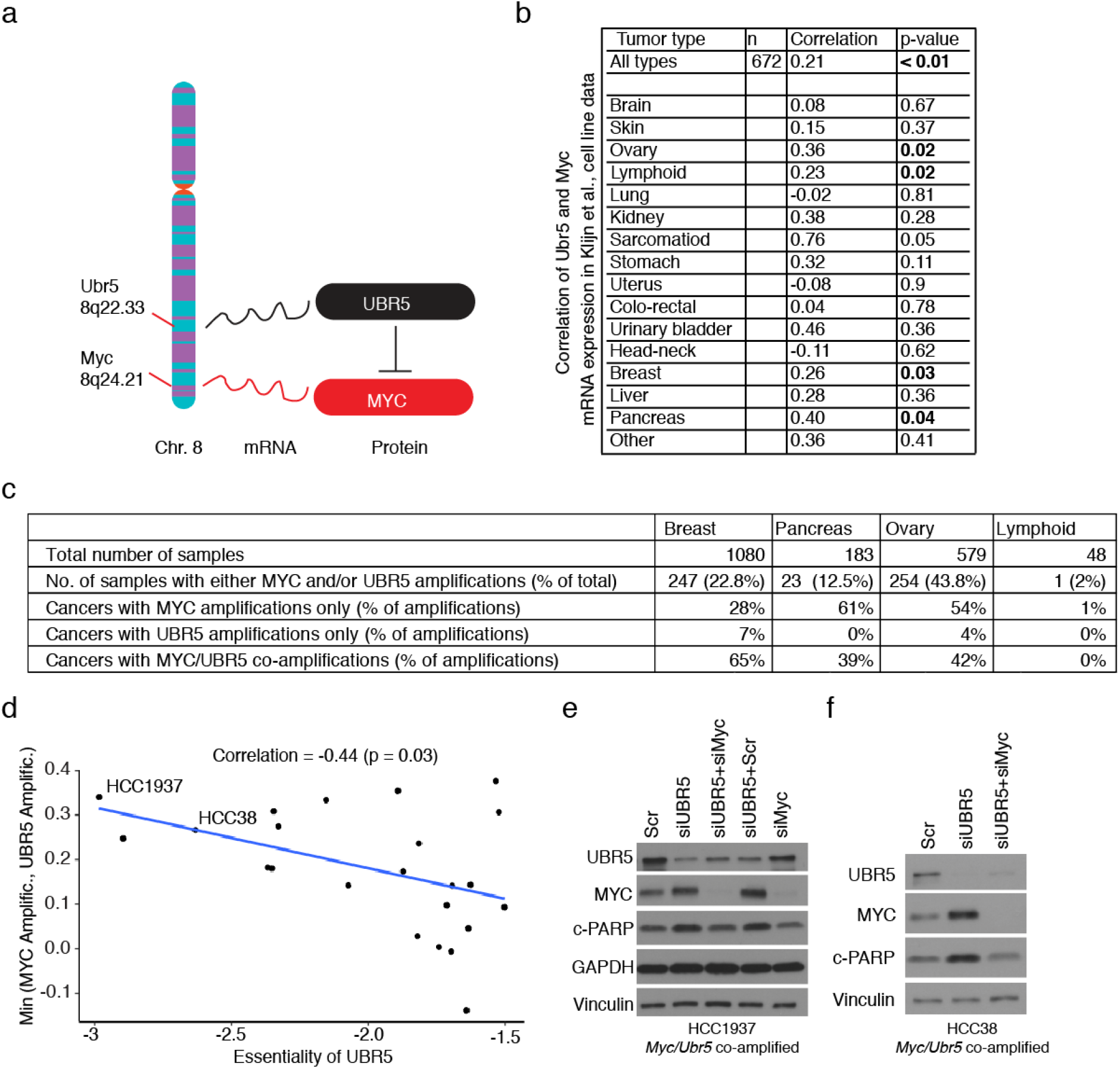
*Ubr5* and *Myc* are co-amplified in human breast cancer. **a**, Schematic figure of Ubr5 and Myc gene locus in the long arm of chromosome 8 and indicated relationship between their expression at mRNA and protein levels. Whereas mRNA expression is predicted to correlate in co-amplified samples, UBR5 is predicted to suppress MYC protein expression in these samples. **b**, Pearson correlation for the relationship between of MYC and UBR5 mRNA expression in cell lines from indicated tumour types. **c**, Percentual distribution of amplification rates of *Myc* and *Ubr5* in TCGA patient samples from the 4 cancer types that displayed significant *Myc* and UBR5 mRNA co-expression in panel (b). **d**, Scatter plot showing the relationship between Ubr5/Myc co-amplification and Ubr5 gene essentiality in the breast cancer cell lines. The minimum of Ubr5 and Myc amplification levels were correlated against the zGARP scores, a measure of UBR5 essentiality in a shRNA dropout screen. **e-f**, Ubr5/Myc co-amplified HCC1937 or HCC38 cells were transfected with indicated siRNAs. Total cell lysates were subjected to immunoblotting with indicated antibodies.

Both *Ubr5* and *Myc* genes are located in the long arm of chromosome 8, suggesting an interesting model where *Ubr5* co-amplification with *Myc* might provide means to control MYC protein levels, and thereby protect cells from apoptosis priming (Fig. 5a). Examination of TCGA cancer patient amplification data for *Ubr5* and *Myc* in the four cancer types with an evidence for mRNA co-expression (Fig. 5b), revealed that the percentage of patient samples with either *Ubr5* and/or *Myc* amplification ranged from 2 % in lymphoid cancers, to 44 % in ovary cancers (Fig. 5c). Interestingly, co-amplification of *Ubr5* and *Myc* was observed in more than 38% of breast, pancreas and ovary cancers, whereas amplification of *Ubr5* alone was a very rare event in these cancer types. In breast cancers, the percentage of cancers with co-amplification exceeded the percentage of cancers with *Myc* amplification alone (Fig. 5c), indicating that *Ubr5* co-amplification and UBR5 protein expression may provide survival benefit for breast cancer cells with *Myc* amplification.

To assess functional relevance of UBR5 in co-amplified breast cancer cells, we correlated UBR5 essentiality index (zGARP score) from a recently published breast cancer cell line drop-out screen (Marcotte, Sayad et al., 2016), with *Myc/Ubr5* gene copy numbers in these same cells. In cell lines with zGARP score < −1.5 for UBR5 (indicating essentiality), we observed a statistically significant correlation between the zGARP score and *Myc/Ubr5* gene copy numbers (Fig. 5d). The functional relevance of UBR5 in co-amplified breast cancer cell lines was further tested in HCC1937 and HCC38 that were among the most UBR5-dependent breast cancer cell lines in the drop-out screen (Marcotte et al., 2016) (Fig. 5d). In support of functional relevance of co-amplification, we observed that UBR5 depletion in these cells did induce MYC protein expression and MYC-dependent apoptosis priming (as observed by PARP cleavage) (Fig. 5e,f).

### UBR5 suppresses drug-induced and MYC-dependent cell killing in *Ubr5/Myc* co-amplified breast cancer cells

Next, we examined TCGA breast cancer patient data in order to study the association between patient survival and amplification status of *Myc* and *Ubr5*. Patients with co-amplification had significantly poorer overall survival than patients without neither of the genes amplified (Fig. 6a). However, as survival of patients with *Myc* amplification alone did not statistically differ from survival of patients with co-amplification, we conclude that potential relevance of UBR5 mediated MYC suppression in co-amplified breast cancers could rather associate with the resistance to therapy-induced apoptosis as was observed in HeLa cells (Fig. 4). To challenge this hypothesis, UBR5 was depleted from HCC38 cells with *Myc/Ubr5* co-amplification, and the cells were treated with FDA approved topoisomerase I inhibitors Irinotecan and Topotecan. Fully supporting the hypothesis, UBR5 depletion resulted in modest MYC-dependent inhibition of colony growth but led to a very potent MYC-dependent sensitisation of cell killing to both topoisomerase I inhibitors (Fig. 6b; the full concentration-dependent dose-response curves are show in Figure S5a-c). We also confirmed a significant, but less pronounced, MYC-dependent priming to Taxol-induced cell killing by UBR5 inhibition in HCC38 cells (Fig. 6c).

**Fig. 6.**
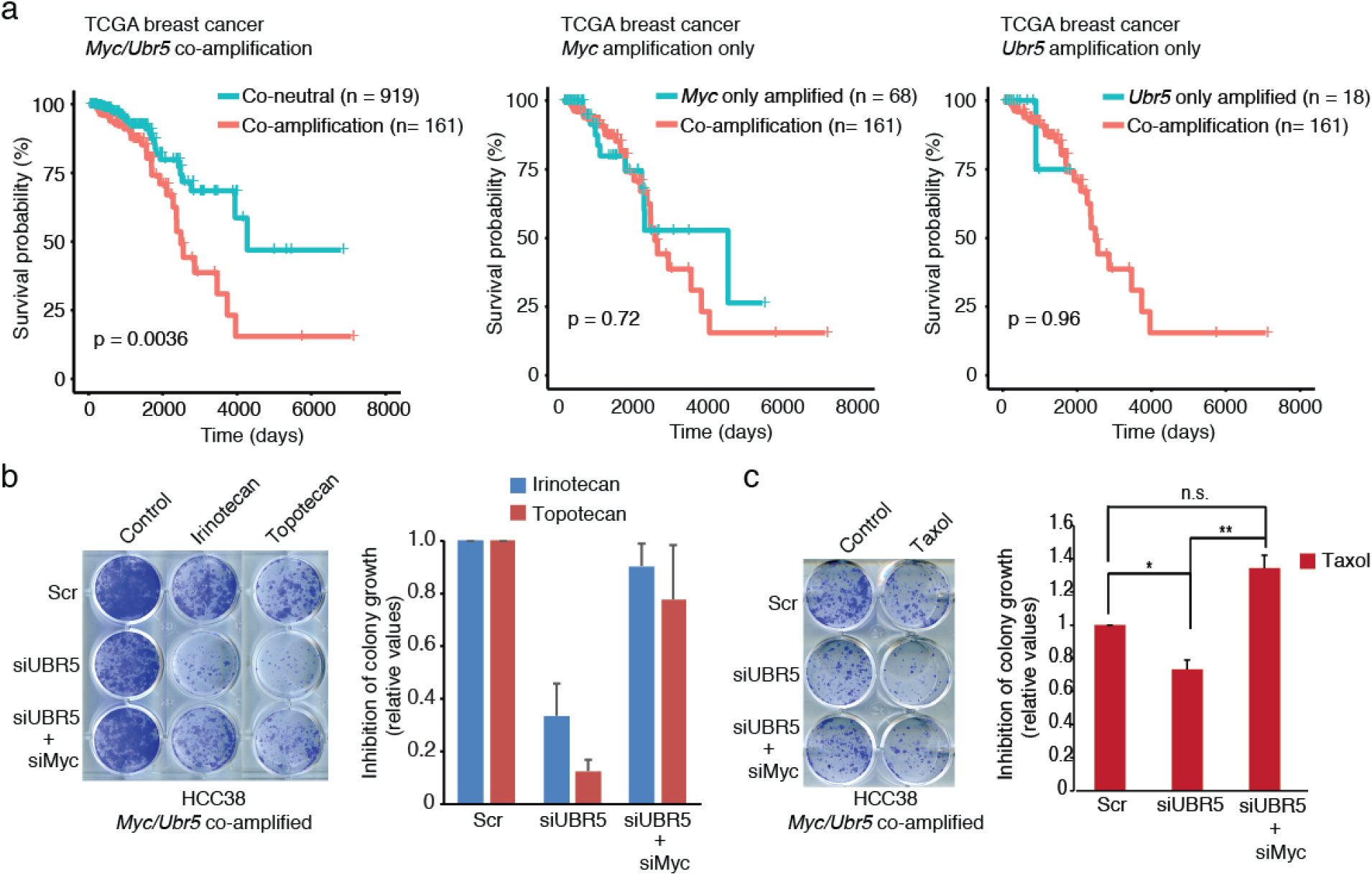
Inhibition of UBR5 sensitizes Ubr5/Myc co-amplified breast cancer cells to drug treatments. **a**, Overall survival analysis of TCGA breast cancer patients that harbour different amplification status of Myc and Ubr5. **b**, Colony growth assay of siRNA transfected HCC38 cells was performed following 24 hours treatment with Irinotecan (80nM) or Topotecan (40nM). Quantification of drug response fold changes relative to Scr siRNA was shown. Error bars show SEM of 2 independent experiments. **c**, Colony growth assay of siRNA transfected HCC38 cells was performed following 24 hours treatment with Taxol (8nm). Quantitation of drug response fold changes relative to Scr siRNA was analysed. Error bars show SEM of 3 independent experiments. *, P < 0.05, **, P < 0.01 by student’s t-test.

Together these results indicate that co-amplification of *Myc* and *Ubr5* might be particularly relevant to breast cancer therapy resistance and that UBR5 inhibition in co-amplified cancers might provide a new therapeutic opportunity in combination setting.

### UBR5 dominates MYC protein expression in individual breast cancer cells *in vivo*

Finally, we examined the *in vivo* relevance of UBR5-mediated MYC protein suppression in human breast cancer tissue samples. To select the most relevant breast cancer subtype for the analysis, the TCGA data was analysed for amplification status of *Myc* and *Ubr5* between different breast cancer subtypes. The results show that there are major differences between breast cancer subtypes in regard to co-amplification frequency. Consistently with published results (Dillon, Mockus et al., 2016), *Myc* amplification was most frequent in basal-type breast cancers and this was also the subtype where co-amplification was most pronounced (Fig. S6a).

Based on this information, we used tissue microarray of 345 samples from 197 human basal-type breast cancers to examine correlation between UBR5 and MYC protein expression *in vivo*. The IHC staining was optimized such that intensities from 0 to +++ were reliably observed with both UBR5 and MYC antibodies (Fig. S6b). Consistently with the hypothesis of UBR5 dominating MYC expression, the percentage of staining positive cells (+, ++, or +++) was greater for UBR5 than for MYC in 78% of the samples (Fig. S6b,c). This finding was re-confirmed by using an independent set of human breast cancer samples (n = 74), and a different staining protocol in an independent laboratory (Fig. S6d).

To validate the UBR5 dominance over MYC protein expression at single cell level, we developed a double immunofluorescence (IF) staining protocol. Antibody specificity was demonstrated by siRNA-mediated depletion (Fig. 1b and S1d), and by secondary antibody only control stainings (Fig. S6e). The double-IF was optimized so that maximal staining intensities were practically indistinguishable with both UBR5 and MYC antibodies (Fig. S6f). Thus, individual cells could be categorized into MYC^high^/UBR5^low^ (green), UBR5^high^/MYC^low^ (red) and MYC^high^/UBR5^high^ (yellow) phenotypes (Fig. 7a). Staining of TMA cores revealed that basal-type breast cancers could be broadly divided into either UBR5 dominant or MYC dominant, or to cancers that showed great heterogeneity at single cell level in regard to UBR5 or MYC dominance (Fig. 7b,c). However, even in those cancers in which the overall staining pattern was heterogeneous (i.e. not predominantly red or green) (Fig. 7b, last column), most of the individual cells were found to be either UBR5 or MYC dominant (Fig. 7c). Quantitative comparison of MYC and UBR5 protein expression at the single cell level from a total of 18 667 cells confirmed both UBR5 dominance, and mutual exclusivity of maximal UBR5 or MYC expression intensity in individual cells (Fig. 7d,e). In this analysis, only 9% of the cells displayed equal UBR5 and MYC protein expression intensities (yellow), whereas UBR5 expression dominated over MYC expression in a majority of individual breast cancer cells (64% vs. 27%)(Fig. 7e). In further support of UBR5 dominance over MYC protein levels in individual basal type breast cancer cells, we re-examined the IHC analysis of breast tumors (Fig. S6g) which was done by using adjacent 3 μM tissue sections enabling us to identify the same cells stained with both UBR5 and MYC antibodies. By these means we could identify several tumour areas where the same individual cells showed a mutual exclusive pattern for maximal staining intensity, i.e. all cells with more intense MYC staining had weaker UBR5 positivity, and vice versa. (Fig. S6g; regions 1 and 2).

**Fig. 7.**
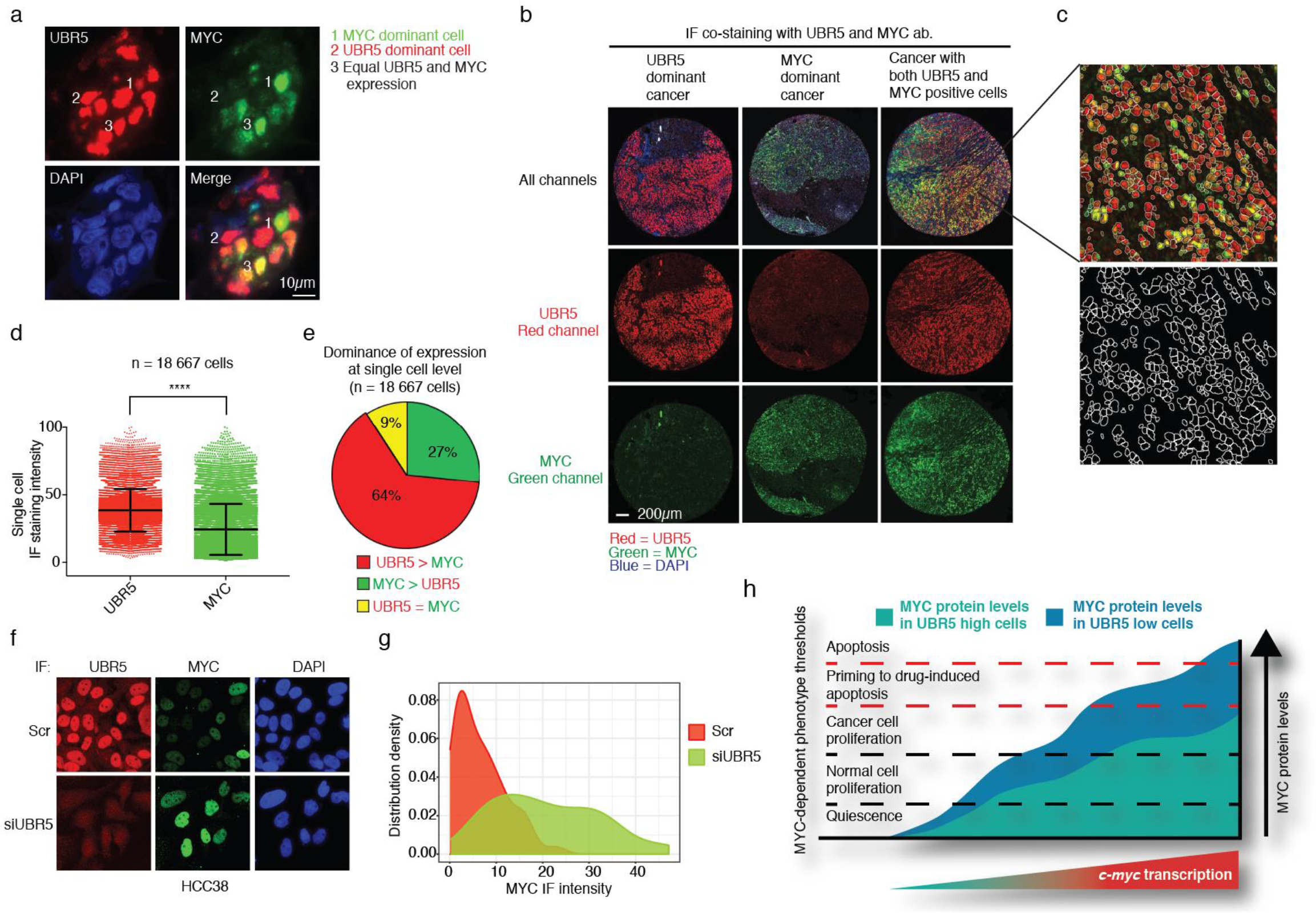
UBR5 dominates MYC protein expression at single cell level in basal type breast cancer *in vivo*. **a**, Intensity of MYC and UBR5 protein from dual immunouorescence staining was used to categorize individual cells into different phenotypes in breast cancer tissues. **b**, Dual immunofluorescent staining of UBR5 and MYC in TMA of human basal-type breast cancer. **c**, The individual cells from dual immunofluorescent staining were gated for quantification by software FIJI. **d-e**, Quantification of UBR5 and MYC staining at single cell level from 18 667 cells by software FIJI. Error bars mean SD. ****, P < 0.0001. **f**, Representative immunofluorescent staining of UBR5 and MYC in HCC38 cells transfected with scrambled (Scr) or UBR5 siRNA. **g**, Quantification of MYC intensity distributions in individual cells from (f). Approximately 50 cells from both scrambled and UBR5 siRNA transfected cells were gated for quantification by software FIJI, and MYC IF staining intensity was blotted in X-axis, whereas the distribution density is blotted on Y-axis. **h**, Schematic presentation of UBR5-mediated control of MYC thresholds in proliferation and apoptosis regulation. MYC protein levels accumulate differently between UBR5 high and low expressing cells upon increased c-myc mRNA expression that in turn can either be due to increased transcription, or by genomic amplification. In UBR5 low expressing cells (blue area), MYC protein accumulates efficiently due to high protein stability, and high MYC protein expressing cells exceed the apoptosis priming threshold. In UBR5 high expressing cells (green area), UBR5 restrains MYC protein accumulation to the levels that maximally support proliferation, but do not exceed the threshold for apoptosis priming even with maximal c-myc mRNA transcription rates.

Taking into an account the finding that some breast cancers displayed heterogeneity in regards to MYC protein expression at single cell level, and this was inversely correlated with high UBR5 expression, we eventually wanted to assess whether UBR5 inhibition could be used to increase the frequency of breast cancer cells with high MYC intensity. Indeed, whereas in scrambled siRNA transfected cell populations only few individual cells expressed MYC at high levels, UBR-depletion greatly increased the number of individual MYC^high^ cells (Fig. 7f,g).

Together these results validate the *in vivo* relevance of the discovered UBR5-mediated suppression of MYC protein expression in human basal type breast cancer cells. The results also reveal a potential novel classification of basal type breast cancers based on UBR5/MYC expression balance at the single cell level.

## Discussion

UBR5 regulates various cellular processes (Gudjonsson, Altmeyer et al., 2012, Sanchez, De Vivo et al., 2016, Shearer et al., 2015, Su, Meng et al., 2011), but the role for UBR5 in both inhibiting and promoting organismal growth has been enigmatic (Callaghan, Russell et al., 1998, Clancy et al., 2003, Kinsella, Dora et al., 2016, Liao et al., 2017, O’Brien et al., 2008, Sanchez et al., 2016, Shearer et al., 2015). The newly identified role of MYC as a critical UBR5 target may provide the thus far missing molecular explanation for the dichotomy. The reported suppression of *Drosophila* tissue growth in *hyd* mutant flies via dMYC is fully consistent with the reported growth suppressor role of Hyd (Callaghan et al., 1998, Shearer et al., 2015). On the other hand, UBR5-mediated suppression of MYC-induced apoptosis observed in cancer cell lines is well in accordance with previously reported oncogenic activities of UBR5 (Clancy et al., 2003, Liao et al., 2017). In support of *in vivo* relevance of these observations, we demonstrate that in most of the basal type breast tumor cells, UBR5 dominates MYC protein expression at the single cell level (Fig. 7d,e). Collectively, we conclude from these results that at least in breast cancers UBR5 finetunes MYC protein expression to the levels that maximally support proliferation, but do not yet prime cells for apoptosis induction (Fig. 7h). Although the conclusions that lower levels of MYC would be more beneficial for the tumour are somewhat counter intuitive, they are fully in accordance with previous results using transgenic MYC mouse models. These studies have demonstrated the importance of MYC thresholds in controlling the delicate balance between proliferation and apoptosis (McMahon, 2014, Murphy et al., 2008, Pelengaris et al., 2002). Although these previous studies convincingly demonstrated that dose of the MYC protein is critical to sustain maximal fitness of the cell, thus far our understanding of the endogenous mechanisms that would control MYC protein expression thresholds have been very limited. Thereby, identification of candidate rheostat proteins such as UBR5, that define threshold levels for MYC-mediated proliferation and apoptosis, may help in future to better understand the functional outcomes of MYC gene regulation.

Many MYC ubiquitin ligases have been identified (Farrell & Sears, 2014), but at least some of them have relatively context-dependent roles in MYC regulation. In this study, UBR5 was identified, among all ubiquitin ligases, as one of the most potential negative regulator of MYC T58A mutant protein expression (Table S1). Validation experiments demonstrated that UBR5–mediated MYC protein regulation was independent of FBW7 and of MYC mRNA regulation, but dependent on the HECT domain of UBR5, and of proteasomal activity. Thereby, the results identify UBR5 as a novel MYC ubiquitin ligase that at least in HeLa cells functions independently of FBW7. However, we naturally cannot fully exclude that the effect of UBR5 on MYC could not be mediated by for example previously identified effect of UBR5 on miRNA regulation (Su et al., 2011). However, UBR5-mediated miRNA regulation was shown to be dependent on PABC, but not on the HECT domain of UBR5 (Su et al., 2011). Notably, in addition to demonstrating both the *in vitro* and the *in vivo* relevance of UBR5-MYC axis, we also show that opposite to some of the other UBR5 functions that are strictly restricted to certain cell types (Kinsella et al., 2016, Saunders et al., 2004), we failed to identify a cell type in which UBR5 inhibition would have not increased MYC protein expression. Therefore, we anticipate these findings to have wide-reaching implications in different biological systems.

Our results indicate *Ubr5* amplification as a novel potential co-dependency with *Myc* amplification in human breast cancer. Based on amplification frequencies of *Ubr5* and *Myc*, it is clear that *Ubr5* amplification alone do not provide significant benefit for breast cancer cells. However, *Myc* amplification without *Ubr5* amplification was also only seen in 28% of the samples with any amplifications, whereas co-amplification was observed in 70% of all cases in which *Myc* was amplified (Fig. 5c). Functionally we show that *Myc/Ubr5* co-amplification frequency correlates with essentially of UBR5 in breast cancer cells (Fig.5d), and that UBR5 inhibition in *Myc/Ubr5* co-amplified breast cancer cells induces MYC-dependent sensitization to cancer drugs (Fig. 6b,c). These results taken together indicate that *Ubr5* co-amplification is useful for a breast cancer cells with *Myc* amplification. Further, these results indicate that pharmacological UBR5 inhibition might sensitize to topoisomerase 1 inhibitors and taxanes in up to 15% (65% of 22.8%; Fig. 5c) of all breast cancer patients stratified based on *Myc/Ubr5* co-amplification. Intriguingly, opposite to breast cancers, Ubr5 amplifications were not observed in lymphoid cancers. This may indicate that UBR5-MYC axis functions differently in lymphoid and solid cancers and that lymphoid cancers might survive with higher MYC levels. The discrepancy between solid and lymphoid cancers in regards to their UBR5 status is also evident by the fact that frequent *Ubr5* mutations (which are predicted to disrupt E3 ligase activity) are observed in mantle cell lymphoma (Meissner, Kridel et al., 2013), which is a cancer type where MYC has also been shown to play an important role.

Results of the study may also have important ramifications to understanding of the functional relevance of MYC protein expression for tumour heterogeneity. We conclude that most of the individual cancer cells in breast tumour tissues harbour MYC levels that efficiently support their proliferation, but do not yet prime them efficiently to apoptosis induction. It is likely that the intratumoural heterogeneity in MYC protein expression at single cell level also translates to incomplete tumor regression in response to various therapies. In support to this, we show that increased MYC positivity in a cancer cell population correlates with degree of cell killing by several cancer therapies. Even though the study did not directly address potential role of UBR5 as a cancer therapy target, our results clearly proposes for a strategy to unleash MYC’s pro-apoptotic activity for cancer therapy. Based on our data, we envision that UBR5 inhibition could harmonize the tumour cells to more uniformly express MYC protein at the levels that would predispose the cells to cell killing by cancer drugs. Optimally this could prevent re-appearance of those cancer cell populations that were protected from the initial drug-induced apoptosis due to their low MYC expression. In addition to the presented data, this model is supported by recent demonstration of MYC as a major *in vivo* determinant of taxane response (Topham et al., 2015), and the data that pretreatment mitochondrial priming correlates with clinical response to cytotoxic chemotherapy in cancer (Ni Chonghaile, Sarosiek et al., 2011).

In summary, this study characterizes a novel mechanism to control MYC protein levels, and thresholds for MYC-induced growth and apoptosis. Our results also provide both a novel molecular explanation for both the tumour suppressive, as well as tumour promoting roles of UBR5 (Liao et al., 2017, O’Brien et al., 2008, Shearer et al., 2015); and support development of specific UBR5 HECT domain inhibitors for cancer combination therapy. Together with recent evidence that both *in vitro* and *in vivo* activity of MYC is highly dependent on post-translational regulation by phosphorylation (Junttila, Puustinen et al., 2007, Myant et al., 2015, Niemela, Kauko et al., 2012), identification of ubiquitination by UBR5 as a critical regulator of MYC-dependent phenotypes clearly further emphasizes the importance of post-translational regulation as a determinant of MYC’s biological activity. In general, better understanding of post-translational regulation of MYC, and of other cancer proteins, might help to explain the overall poor concordance between genomic alterations and corresponding protein activities in cancer (Mertins et al., 2016).

## Acknowledgements

The authors thank Prof. Johanna Ivaska and Prof. Lea Sistonen for valuable comments to the manuscript. The authors thank for Prof. Dean Felsher providing osteosarcoma-MYC-off cell line. Prof. Darren Saunders and Charles Watts are thanked for UBR5 plasmids, and Prof. Bruno Amati and Rosalie Sear for MYC plasmids. Taina Kalevo-Mattila is thanked for excellent technical help, and Jouko Sandholm at the Turku Centre for Biotechnology Cell Imaging Core facility for help with imaging. Sinikka Collanus is thanked for her help in IF and IHC. Samu Kurki, Eliisa Löyttyniemi and Srikar Nagelli are acknowledged for providing us help for statistical analysis of IHC data. Birgit Samans is thanked for the support with the statistical analysis of the high-content siRNA screen image assay. Finnish Functional Genomics Centre and Bioinformatics Unit of the Turku Centre for Biotechnology provided service for RNA-sequencing analysis. This study was supported by funding from Sigrid Juselius Foundation, Finnish Cancer Association, Biocenter Finland, Deutsche Forschungsgemeinschaft grant (Ei222/12-1), Foundation Martin Escudero, Varsinais-Suomi Regional Fund, and Instrumentarium Science Foundation.

## Materials and methods

### Cell culture and transfection

HeLa, U2OS, T98G, MCF10A, HEK293, HCC38, HCC1937, IMR-32 and MDA-MB-231 cell lines were obtained from American Type Culture Collection. Osteosarcoma-MYC-off cell line(Jain et al., 2002) was a generous gift from Professor Dean Felsher (Stanford University). HeLa, U2OS, T98G, HEK293, Osteosarcoma-MYC-off and MDA-MB-231 cell lines were cultured in DMEM (Sigma). HCC38, -HCC1937 and IMR-32 cell lines were cultured in RPMI (ATCC-modified vision, Thermo Fisher Scientific). MCF10A cells were cultured as described previously (Myant et al., 2015). Drosophila S2 cells were cultured in Schneider’s Drosophila Medium (Thermo). All growth mediums were supplemented with 10% heat-inactivated FBS (Gibco), 2 mmol/L L-glutamine, and penicillin (50 units/mL)/streptomycin (50 mg/mL). All cell lines were cultured in a humidified atmosphere of 5% CO_2_ at 37°C. The cells were treated with 20μM MG-132 (474791-5MG, Calbiochem), 6 hours. Cycloheximide, doxorubicin, camptothecin, taxol and irinotecan were from Sigma-Aldrich. Topotecan was from Selleck chemicals.

GFP-UBR5, GFP-UBR5ΔHECT, Flag-UBR5, and Flag-UBR5ΔHECT were kind gifts from Darren Saunders & Charles Watts(Gudjonsson et al., 2012, Henderson, Russell et al., 2002). V5-MYC, V5-MYC^T58A^ and V5-MYC^S62A^ have been described previously (Yeh, Cunningham et al., 2004). HA-ubqiuitin was a kind gift from professor Lea Sistonen (Åbo Akademi University). Plasmids were transfected with Lipofectamine® 2000 Transfection Reagent (Thermo Fisher Scientific) according to the manufacturer’s instructions. After 48 hours transfection, cell lysate was collected. Small interfering RNA (siRNA) transfections were performed with Oligofectamine™ Transfection Reagent (Thermo Fisher Scientific) following to the manufacturer’s protocol. Three days after transfections, cells were harvested for analysis. siRNA target sequences are in table 1.

### siRNA library and high-content screening

siGENOME RNA oligonucleotides were purchased from Thermo Scientific Dharmacon (G-005000, lot-050915), and a siUbiquitin sublibrary containing 591 pooled siRNAs against known and putative E1, E2 and E3 ubiquitin ligases was generated. To reduce potential off-target effects, a pool of four individual siRNAs was transfected for each candidate ubiquitin ligase gene. Clonal selected U2OS cells stably expressing MYCT58A under the control of the retroviral LTR were transfected with 50nM siRNA in a 96-well format using Dharmafect1 (Dharmacon) or Lipofectamine RNAiMAX (Invitrogen). 24 hours after transfection, medium was replaced by fresh culture medium. 24 hours later, to halt Myc mRNA translation and reduce both endogenous MYC WT and ectopically expressed MYCT58A protein levels, 100μgmL–1 cycloheximide was added to the medium for 3,5 hours. Cells were fixed with 4% paraformaldehyde and subjected to indirect immunofluorescence using a MYC antibody (N-262, Santa Cruz Biotechnology) followed by Alexa488-conjugated secondary antibody (Invitrogen). For automated immunofluorescent-based data acquisition, the BD PathwayTM 855 bioimager (BD Biosciences) was used. Cellular MYC protein levels were quantified with the BD AttoVisionTM software (BD Biosciences). Data quality was assessed by z-factor analysis. Relative changes in MYC protein levels were calculated using the z-score.

### Colony formation assay

The optimized numbers of cells were seeded in 12-well plates directly or after 24h transfection for siRNA transfected cells until formation of colonies. After 24 hours seeding, colonies were treated with indicated concentration of chemical drugs for another 24 hours. Cell colonies were fixed with cold methanol and stained with 0.2% crystal violet solution (made in 10% ethanol) for 15 minutes at room temperature each. Excess stain was removed by repeated washing with PBS. Plates were dried and scanned with Epson perfection V700 scanner. Quantifications were performed with ColonyArea ImageJ plugin(Guzman, Bagga et al., 2014) and graphs were plotted using the area % values.

### Immunoblotting and immunoprecipitation

Cultured, siRNA transfected and/or treated cells were lysed in RIPA buffer (50 mM Tris-HCl pH 7.5, 0.5 % DOC, 0.1 % SDS, 1% NP-40, and 150mM NaCl) with protease and phosphatase inhibitors (4693159001 and 4906837001, Roche). The lysate was sonicated, added with 6X SDS loading buffe, boiled and resolved by 4-20% precast protein gels (456-1093 and 456-1096, Biorad). Proteins were transferred to PVDF membranes (1704156, Biorad). Membranes were blocked in 5% Milk-TBS-Tween 20 for 30 minutes under RT, and then incubated with primary antibodies overnight at 4°C. Secondary antibodies was incubated in 5% Milk-TBS-Tween 20 for 1 hour under RT, and developed by ECL western blotting substrate (32106, Pierce). The following are antibodies used for western blot: UBR5 (sc-515494, Santa cruz), MYC (ab32072, Abcam), phospho-S62MYC (ab78318, Abcam), cleaved PARP (ab32064, Abcam), BIM (2933, Cell signaling), Vinculin (sc-25336, Santa cruz), GAPDH (5G4-6C5, HyTest Ltd), HA (3724, Cell signaling), Flag (F3165, Sigma), GFP (sc-9996, Santa cruz), V5 (R960-25, Invitrogen), Histone H3 (ab1791, Abcam) and Lys48-Specific antibody (05-1307, Millipore). Secondary antibodies are from Dako (P0447 and P0399). Densitometric analysis of the blots was performed using ImageJ. For immunoprecipitation, the cells were lysed in IP buffer (150mM NaCl, 1% NP-40, 50mM Tris pH 8.0) supplemented with protease and phosphatase inhibitors. Cell lysates were centrifuged and immunoprecipitated with the indicated antibodies for 2 h at 4 °C, following by adding Protein A-Sepharose or protein G-Sepharose beads (Sigma) overnight. The beads were washed with IP buffer, boiled in SDS sample buffer and analysed by immunoblotting. MYC (N-262, sc-764), and V5 agarose (A7345, Sigma) were used for immunoprecipitation.

For double-stranded RNA (dsRNA) mediated RNAi in Drosophila S2 cell, the DNA fragment of hyd was amplified with primers flanked by T7 promoter, and dsRNA was produced using the TranscriptAid T7 High yield Transcription Kit (Thermo). To knock down Hyd, S2 cells were cultured with 5 ug/ml dsRNA for 120h. After cultured with dsRNA, S2 cells were washed with PBS, and lysed with lysis buffer (50 mM Tris-HCl pH 7.8, 150 mM NaCl, 0.5% NP-40) supplemented with Pierce protease inhibitors (Thermo) and Pierce phosphatase inhibitor (Thermo). Equal protein levels were confirmed by a control western blot of rabbit anti-Kinesin (Cytoskeleton) from the cleared lysates. To immunoprecipitate dMyc, lysates were incubated 2 hours with anti-dMyc beads. Anti-dMyc beads were made from rabbit anti-dMyc antibody (Santa Cruz) and 50% protein A Sepharose beads (Amersham). Immunoprecipitates were washed five times with NP-40 lysis buffer, boiled in 2X SDS sample buffer, and resolved on 8% SDS-PAGE for analysing via Western blotting. Immunoprecipitated Myc was detected using mouse anti-Myc antibody (a kind gift from Dr. Peter Gallant).

### Immunofluorescence staining

Cells plated on chambered coverslip (80826, Ibidi) were transfected with scrambled siRNA or UBR5 siRNA. After 72 hours transfection, the cells were fixed with 4% paraformaldehyde 15 minutes under room temperature, and then cells were permeabilized with 0.5% Triton X-100 in PBS on ice for 5 minutes. Next, the cells were blocked by 10% normal goat serum (ab7481, Abcam) diluted in PBS for 30 minutes, and followed by incubating the primary antibodies anti-UBR5 (sc-515494, Santa cruz) and anti-MYC(ab32072, Abcam) overnight at 4°C. Subsequently, cells were washed with PBS and incubated with secondary antibodies, Alex Fluor 594 goat anti-Mouse IgG (A-11005, Invitrogen) and Alex Fluor 488 goat anti-rabbit IgG (A-11008, Invitrogen) for 1 hour under room temperature. After secondary antibody incubation, the cells were washed with PBS and nuclei were stained with DAPI (D1306, Invitrogen) in PBS at RT for 10 min. Images were acquired with confocal microscope (LSM780, Carl Zeiss).

### Proximity ligation assay

The PLA assay was performed according to the manufacturer’s protocol (Duolink® PLA, Sigma). Briefly, HeLa cells plated on coverslips were transfected with scrambled siRNA or MYC siRNA. After 72 hours transfection, the cells were fixed with 4% paraformaldehyde 15 minutes under room temperature, and then cells were permeabilized with 0.5% Triton X-100 in PBS on ice for 5 minutes. Next, cells were blocked with blocking solution, and incubated in a pre-heated humidity chamber for 30 min at 37°C, followed by incubating the primary antibodies (in blocking solution) anti-UBR5 (sc-515494, Santa cruz) and anti-MYC (ab32072, Abcam) overnight at 4°C. Subsequently, cells were washed with buffer A, and the PLA probe was incubated in a preheated humidity chamber for 1 hr at 37°C, followed by ligase reaction in a preheated humidity chamber for 30 minutes at 37°C. Next, amplification polymerase solution for PLA was added, followed by incubating the cells in a pre-heated humidity chamber for 100 min at 37°C. After amplification, the coverslips were washed with buffer B, and mounted with DAPI. PLA signal was detected by using a confocal microscope (LSM780, Carl Zeiss).

### Caspase3/7 activity assay

Caspase3/7 activity was measured by luminescence-based method, which utilize a substrate containing Caspase3 and Caspase7 target peptide DEVD, named Caspase-Glo® 3/7 assay (G8091, Promega). The assay was performed following the manufacturer’s protocol in white polystyrene 96-well plates (Nunc, Thermo Fisher Scientific Inc.) and luminescence was measured with Perkin Elmer Victor-2 Plate Reader (PerkinElmer Inc.).

### Quantitative RT-PCR and RNA sequencing

Total RNA from HeLa was isolated from by using RNAeasy kit (Qiagen) according to the manufacturer’s protocol. cDNA was synthetized using 1 μg of DNAse I (Life Technologies) treated RNA using MMLV RNase H minus reverse transcriptase (Promega) and random hexamer primers (Promega).

Real-time PCR analysis of cDNA samples was performed with specific primers and probes designed by using Assay Design Center (Roche). Primers and probes used here are available upon request. Expression of each gene was presented as the percentage of interested mRNA expression relative to the control gene expression.

RNA was extracted from S2 cells using Nucleospin RNA II kit (Macherey-Nagel) and cDNA was synthesized using SensiFAST cDNA Synthesis kit (Bioline) according to the manufacturer’s protocol. qPCR was performed with Light cycler 480 Real-Time PCR System (Roche) using SensiFAST SYBR No-ROX Kit (Bioline).

RNA-seq was performed with RNAs derived from HeLa cells transfected with indicated siRNAs by using Illumina RNA-sequencing at Finnish Functional Genomics Centre (Turku, Finland). The samples were sequenced with the HiSeq2500 instrument using single-end sequencing chemistry with 50 bp read length.

### *Drosophila* genetics

Fly stocks used in this study are: Myc RNAi (VDRC 2947), hyd mutant allele K3.5(Lee, Amanai et al., 2002) and MARCM82b (a kind gift from Dr. Osamu Shimmi). To induce MARCM clones in developing wing discs(Lee & Luo, 1999), larvae were heat-shocked for one hour at 37°C at 72 hours after egg laying. Wing imaginal discs were dissected from late third instar larvae, fixed in 4% formaldehyde for 30 min, after washing in PBT (0.3% Triton X 100 in PBS), samples were mounted with Vectashield Mounting Medium with DAPI (Mediq), and imaged using a Zeiss LSM 700 microscope. Clone roundness was analyzed as described earlier (Prober & Edgar, 2002). Volume per cell and pH3 positive cells were counted using Imaris software. To induce MARCM clones in larvae fat body, larvae were heat-shocked for one hour at 37°C at 24 hours after egg laying. Fat bodies were dissected from third instar larvae, fixed in 4% formaldehyde for 30 min, and washed in PBT (0.3% Triton X 100 in PBS). After blocking in 5% BSA in PBT for 3 hours at RT, primary antibody (anti-FBL, 1:500, Abcam) was incubated at 4 C o/n. Primary antibody were washed with PBT and secondary antibody (anti-mouse Alexa fluor 647, Life Technologies) was incubated for 4 hours at RT. After three washes, samples were mounted with Vectashield Mounting Medium with DAPI (Mediq), and imaged using a Zeiss LSM 700 microscope.

Full genotypes in Fig 3d:

yw, hsFLP/+; Tub-G4, UAS-mCD8 GFP/+; FRT82b, Tub-Gal80/FRT82, hydK3.5

Full genotypes in Fig 3e and f:

<u>Ctrl</u>: yw, hsFLP/+; Tub-G4, UAS-mCD8 GFP/+; FRT82b, Tub-Gal80/FRT82b
<u>Myc RNAi</u>: yw, hsFLP/+; Tub-G4, UAS-mCD8 GFP/Myc RNAi; FRT82b, Tub-Gal80/FRT82b
<u>hyd mutant</u>: yw, hsFLP/+; Tub-G4, UAS-mCD8 GFP/+; FRT82b, Tub-Gal80/FRT82, hyd K3.5
<u>hyd mutant, Myc RNAi</u>: yw, hsFLP/+; Tub-G4, UAS-mCD8 GFP/Myc RNAi; FRT82b, Tub-Gal80/ FRT82b, hyd K3.5

### Breast cancer IHC and IF analysis

Intrinsic classification and identification of basal differentiation in TNBC were performed based on detection of ER, PR, Ki-67, CK5/6 and EGFR IHC, and HER2/Chr17 double ISH using automated immunostaining BenchMark XT machine (Roche Diagnostics / Ventana) (Lakhani SR, Ellis IO, Schnitt SJ, Tan PH, van de Vijver MJ (eds.) (2012) WHO Classification of Tumors of the Breast pp 10–11. IARC: Lyon).

Tissue material was treated according to standard histology practice, i.e. fixed in buffered formalin (pH 7.0) and embedded into paraffin blocks. Tissue microarrays (TMAs) were prepared by collecting two 1.5 mm diameter tissue cores from the representative tumor area, defined by an experienced breast cancer pathologist (PK), of each breast cancer patient. Both immunohistochemistry (IHC) and double immunofluorescence (IF) were performed on sections cut at 3.5 μm. For MYC (ab32072, Abcam, Y69), IHC was performed with Lab Vision Autostainer 480 (Thermo-Fisher Scientific, Fremont, CA, USA) and detected with PowerVision+ polymer kit (DPVB+110HRP; Immunovision Technologies, Vision Biosystems, Norwell, MA, USA) according to standard protocol with diaminobenzidine as chromogen. Before staining, tissue sections were deparaffinized and treated twice for 7 min each in Target Retrieval Solution, pH 9 (S2367, Dako, Glostrup, Denmark) in a microwave oven for antigen retrieval. MYC antibody was applied at a dilution of 1:250. An automated immunostaining machine Discovery XT (Roche Diagnostics/Ventana Medical Systems, Tucson, AZ, USA) was used for UBR5 (sc-515494, Santa Cruz Biotechnology) IHC and UBR5/MYC double IF. Deparaffinization, epitope retrieval (standard option, Cell Conditioning 1 reagent, 950-124, Roche/Ventana) and primary antibody incubation (40 min at 37° C, dilution 1:500 for UBR5-IHC, and 1:1000 and 1:50 for UBR5 and MYC, respectively, for double-IF) were done on the platform. OmniMap HRP (760-4310, Roche/Ventana) and ChromoMap DAB Kit (760-159, Roche/Ventana) were applied for detection of UBR5 IHC. For double IF, OmniMap HRP (Roche/Ventana, 760-4310 and 760-4311 for antimouse and anti-rabbit, respectively) together with Rhodamine and FAM fluorescent substrates (Roche/Ventana, 760-233 and 760-243, respectively) were used. Finally, IF slides were mounted applying ProLong® Gold antifade reagent with DAPI (P36935, Molecular Probes by LifeTechnologies). Two slides were stained as controls for double IF. One, by replacing UBR5 primary antibody, and the other, by replacing MYC primary antibody with antibody diluent.

### Analysis of mutual exclusivity of MYC and UBR5 on single cell level

To analyze MYC and UBR5 expression levels on a single cell level, we generated an automated image analysis macro for the open source software FIJI (Schindelin, Arganda-Carreras et al., 2012). This macro was used to analyse cells in 20 histological samples. In brief the macro defines a single cell by creating a mask and applying a colour threshold on a merged (MYC-green and UBR5-red) image after background subtraction and image smoothing. The created mask is further processed by applying a watershed function and followed by a selection of a minimum particle size, to precisely distinguish individual cells. Every separated cell is then given an ID and both signal levels, MYC (green) and UBR5 (red), are measured and displayed.

The measurement output of the macro was further analyzed using Microsoft Excel. First, we normalized the signal intensities in both channels against the maximum green/red signal in the dataset. The difference of these values was calculated and if the signal intensity was more than 10% higher in one channel than in the other, it was defined as dominant, and the cell was marked as “green” or “red”. Differences in signals intensities within the ±10% were considered equal and cells were marked as “yellow”. Finally, the number of “green”, “red” and “yellow” cells was calculated.

### Data analysis and statistical testing

All the data analysis and statistical test were performed in the R statistical programming environment (http://R-project.org). The open-source R scripts and the data files can be accessed from the GitHub repository (https://github.com/abishakGupta/analysisMYC). The specific data analyses are explained below:

#### Expression correlation

The RNA-seq data for the 675 cell lines were extracted from Klijn *et. al.* (Nature Biotech. 2015). The median values of UBR5 and c-MYC RNA expression data were calculated and their association was computed using the Pearson correlation coefficient. The statistical significance (p-values) of the Pearson correlation coefficient levels was assessed with the Fisher’s z-transformation using inbuilt functions in R.

#### Co-amplification analysis and Survival analysis

Copy number calls using the GISTIC 2.0 algorithm were downloaded from cBioPortal (http://www.cbioportal.org/) for the four cancer types of interest: breast, pancreas, ovary and diffuse large B-cell lymphoma as a representative of lymphoid tissue type. In this analysis, each gene has a discretized value denoting its copy number status; −2 for homozygous deletion, −1 for hemizygous deletion, 0 for neutral, 1 for gain, and 2 for high level amplification. Patient samples with copy number status of 2 and 0 were considered as amplified and neutral, respectively, for Ubr5 and Myc separately. The percentage of samples with Ubr5 and Myc co-amplification was computed based on these selected patient samples separately for each cancer type.

Kaplan–Meier analysis and univariate Cox proportional hazard test was used to assess the difference in overall survival between two groups of patients. The groups were defined as co-amplified and co-neutral based on the copy number status of Ubr5 and Myc in the patient tumors.

#### Essentiality analysis

To assess the relationship of the co-amplification with the gene functionality, we extracted the amplification and gene essentiality data of breast cancer cell lines reported in(Marcotte et al., 2016). We used the minimum Ubr5 and Myc amplification values, and computed their association with the gene essentiality, as measured by zGARP scores, using the Pearson correlation coefficient. The statistical significance (p-values) of the Pearson correlation coefficient levels was computed using the Fisher’s z-transformation.

